# Modulating Motor Cortex Plasticity via Cortical and Peripheral Somatosensory Stimulation

**DOI:** 10.1101/2024.12.12.628248

**Authors:** Shancheng Bao, Yuming Lei

**Affiliations:** Program of Motor Neuroscience Department of Kinesiology & Sport Management Texas A&M University, College Station, TX, 77843

**Keywords:** Primary somatosensory cortex, Peripheral nerve, Primary motor cortex, Continuous theta-burst stimulation, Transcutaneous electrical nerve stimulation

## Abstract

The interaction between the motor and somatosensory systems is essential for effective motor control, with evidence indicating that somatosensory stimulation influences the excitability of the primary motor cortex (M1). However, the mechanisms by which repetitive stimulation of both cortical and peripheral somatosensory systems affects M1 plasticity are not well understood. To investigate this, we examined the effects of continuous theta-burst stimulation (cTBS) applied to the primary somatosensory cortex (S1) and transcutaneous electrical nerve stimulation (TENS) of the median nerve on various measures of corticospinal excitability and M1 intracortical circuits. Specifically, we assessed motor-evoked potentials (MEPs), short-latency intracortical inhibition (SICI), intracortical facilitation (ICF), and short-latency afferent inhibition (SAI) before and after administering cTBS and TENS. Our results demonstrated that cTBS increased MEPs for at least 50 minutes, whereas TENS increased MEPs for 10 minutes. Neither cTBS nor TENS had an impact on SICI and ICF. However, cTBS decreased SAI, while TENS did not affect SAI. The sham procedures for both cTBS and TENS did not produce significant changes in MEPs, SICI, ICF, or SAI. These findings suggest that both cortical and peripheral somatosensory stimulation modulate corticospinal excitability, with the effects of cortical stimulation being more prolonged. Neither type of stimulation influences inhibitory and excitatory intracortical neural circuitry within M1. Notably, cortical somatosensory stimulation modulates the interaction between M1 and S1, whereas peripheral somatosensory stimulation does not. This study elucidates distinct mechanisms through which cortical and peripheral somatosensory stimulation influence M1 plasticity.

**NEW & NOTEWORTHY:** This study identifies distinct mechanisms through which cortical and peripheral somatosensory stimulation influence motor cortex (M1) plasticity. Both continuous theta-burst stimulation (cTBS) of the primary somatosensory cortex (S1) and transcutaneous electrical nerve stimulation (TENS) of the median nerve enhance corticospinal excitability, with cTBS exhibiting longer-lasting effects. Importantly, cTBS, but not TENS, modulates the interaction between M1 and S1. These findings form the basis for developing targeted somatosensory interventions aimed at modulating motor function.

## Introduction

The somatosensory system is integral for receiving and processing sensory information from the body and is crucial in executing voluntary movements (Kaas, 1993; Johansson and Flanagan, 2009; Lei et al., 2018). Disruptions within this system, whether cortical or peripheral, can result in significant impairments in movement coordination, precision, and the acquisition of new motor skills (Rothwell et al., 1982; Brochier et al., 1999; Blennerhassett et al., 2007; Kumar et al., 2019). Inhibition of the primary somatosensory cortex (S1) with GABA injections has been shown to impair finger coordination in monkeys (Hikosaka et al., 1985; Brochier et al., 1999). Additionally, lesion or inhibition of S1 disrupts the learning of new motor behaviors in various species, including cats (Sakamoto et al., 1989), mice (Mathis et al., 2017), and monkeys (Pavlides et al., 1993). In humans, inhibition of S1 can interfere with the consolidation of motor memory (Kumar et al., 2019; Wang et al., 2023). Peripheral deafferentation in the hands significantly impairs fine hand movements in monkeys (Mott and Sherrington, 1895), while humans with somatosensory deafferentation exhibit motor impairments (Sainburg et al., 1993, 1995). Individuals with complete somatosensory deafferentation struggle with tasks requiring fine motor skills, such as grasping, writing, or fastening buttons (Rothwell et al., 1982). Conversely, stimulating the somatosensory system can enhance motor function. For example, S1 stimulation significantly improves upper extremity sensorimotor function in stroke patients (Meehan et al., 2011; Brodie et al., 2014; Pundik et al., 2021), and peripheral nerve stimulation of the paretic upper extremity enhances motor performance post-stroke (Celnik et al., 2007; Klaiput and Kitisomprayoonkul, 2009; Knutson et al., 2012). These findings underscore the critical role of the somatosensory system in motor function and emphasize the potential of targeted somatosensory stimulation in modulating motor capabilities.

Targeting the somatosensory system can be achieved through various methods, encompassing both peripheral and cortical somatosensory stimulation. Transcutaneous electrical nerve stimulation (TENS) administers electrical impulses to specific nerves, thereby modulating activity within the somatosensory system. Research has demonstrated that TENS can have a range of effects, including increasing endogenous neurotransmitter levels (Sun et al., 2018; Meyer-Frieem et al., 2019), elevating the N20 component of somatosensory evoked potentials (Ferretti et al., 2007), and enhancing blood oxygen level-dependent (BOLD) effects in S1 (Schabrun et al., 2012). Additionally, techniques such as transcranial magnetic stimulation (TMS) can be employed to directly modify activity in S1. Repetitive TMS (rTMS) has been shown to induce changes in SEP amplitude and improve tactile discrimination learning in rats (Mix et al., 2010; Benali et al., 2011). In human studies, rTMS over S1 has been observed to alter sensorimotor network activity patterns within S1 and M1, with effects persisting for up to 120 minutes following stimulation (Pleger et al., 2006). Neuropsychological studies have also reported improvements in spatial acuity (Ragert et al., 2003), tactile perception (Tegenthoff et al., 2005), and distance discrimination (Pleger et al., 2006) after rTMS application over S1. Despite evidence showing that TENS and TMS can induce neuroplastic changes within the somatosensory system (Ragert et al., 2003; Pleger et al., 2006; Ferretti et al., 2007; Benali et al., 2011; Sun et al., 2018; Meyer-Frieem et al., 2019), the mechanisms by which stimulation of both the cortical and peripheral somatosensory systems modulate M1 plasticity remain largely unknown. Previous studies have yielded contrasting findings regarding the effects of TENS and TMS on M1 excitability. Some studies showed that TENS leads to a suppression of M1 excitability (Maertens et al., 1992; Tokimura et al., 2000), whereas others have reported an enhancement of M1 excitability (Veldman et al., 2015; Ferretti et al., 2007). Similarly, some studies showed that rTMS over S1 can influence corticospinal output to the hand (Jacobs et al., 2014) and affect the inhibitory influence of tactile inputs on M1 excitability (Bao et al., 2024). Conversely, other research has found no significant effects of rTMS applied over S1 on M1 excitability (Ishikawa et al., 2007; Katayama et al., 2010). Here, we examined the effects of cortical and peripheral somatosensory stimulation using cTBS and TENS on M1 plasticity.

This study investigated the effects of cTBS applied to S1 and TENS of the median nerve on various measures of corticospinal excitability and intracortical circuits within M1. In Experiment 1, we assessed motor-evoked potentials (MEPs), which reflect corticospinal excitability from both cortical and subcortical levels (Petersen et al., 2003), before and after the administration of cTBS and TENS. In Experiment 2, we measured short-latency intracortical inhibition (SICI) and intracortical facilitation (ICF), which indicate GABAA receptor activities (Kujirai et al., 1993; Di Lazzaro et al., 2006) and glutamate-driven excitatory functions (Ziemann et al., 1996, 1998) in M1, respectively, before and after the administration of cTBS and TENS. Prior research suggests that these intracortical circuits play a role in M1 cortical plasticity (Kujirai et al., 1993; Ziemann et al., 1996). In Experiment 3, we measured short-latency afferent inhibition (SAI), reflecting S1-M1 interactions characterized by M1 inhibition induced by afferent inputs through paired-pulse stimulation (Tokimura et al., 2000; Lei and Perez, 2017; Davis et al., 2022), before and after the administration of cTBS and TENS. While TENS of the median nerve primarily sends bottom-up ascending projections to the cortex affecting M1 plasticity, cTBS applied to S1 influences M1 plasticity mainly through the connections between S1 and M1. Therefore, we hypothesized that distinct mechanisms exist through which cortical and peripheral somatosensory stimulation influence M1 plasticity.

## Materials and Methods

### Participants

This study comprised seventy-three right-handed, healthy participants aged between 18 and 30. All participants were unfamiliar with the paradigm and the purpose of the study. The Institutional Review Board of Texas A&M University approved all experimental protocols. Furthermore, each participant provided written informed consent prior to participation, in accordance with the Declaration of Helsinki, as approved by the local ethics committee at Texas A&M University.

### Electromyographic (EMG) recordings

Surface EMG was recorded using disposable Ag-AgCl electrodes with a 10 mm diameter, positioned on the skin surface over the muscle belly. The EMG signals were amplified and filtered within a 30–2000 Hz range utilizing a Neurolog System bioamplifier (Digitimer, UK). These signals were then digitized at a sampling rate of 10 kHz using a CED Micro 1401 A/D converter (Cambridge Electronic Design, UK) and subsequently stored on a computer for later analysis.

### Transcranial Magnetic Stimulation (TMS)

TMS was conducted using two systems: the DuoMAG MP-Dual TMS system (Brainbox Ltd., UK) and the Magstim Rapid^2^ system (Magstim, UK). Neurophysiological measurements (MEP, SICI, ICF, and SAI) were obtained via the DuoMAG MP-Dual TMS system, while cTBS was administered using the Magstim Rapid^2^ system. Measurements of MEPs, SICI, ICF, and SAI were recorded from the First Dorsal Interosseous (FDI) muscle of the right hand via EMG using the DuoMAG MP-Dual TMS system. The TMS coil was positioned tangentially on the scalp at a 45° angle from the midline, with its handle oriented laterally and posteriorly to induce a posterior-anterior current in the brain. The optimal stimulation position, known as the hotspot, was identified as the location where the largest MEP in the FDI muscle was evoked with minimal stimulus intensity. This hotspot was marked using a frameless neuronavigation system (Brainsight Ltd., Canada) to ensure consistent coil placement during the study. The resting motor threshold (RMT) was defined as the lowest stimulus intensity that produced MEPs exceeding 50 μV in peak-to-peak amplitude above background EMG activity in at least 5 out of 10 trials. This study administered cTBS to S1 using a 70 mm figure-of-eight coil paired with the Magstim Rapid^2^ system. The stimulation site for S1 was determined by measuring 2 cm posterior to the M1 hotspot, utilizing the neuronavigation system. Previous research has validated this method to identify the homologous effector location in S1 corresponding to M1 (Ishikawa et al., 2007; Conte et al., 2016; Tsang et al., 2014). Stimulation at S1 using this technique has been shown to modulate SEP (Wolters et al., 2005; Ishikawa et al., 2007). Consistent with these findings, our prior studies have demonstrated that cTBS over S1 decreases the amplitude of N20 SEP (Wang et al., 2024), indicating reduced excitability of S1. We administered cTBS at an intensity of 70% RMT, following a protocol consisting of 600 pulses at 30 Hz, with bursts separated by 167 ms intervals (Gentner et al., 2008; Goldsworthy et al., 2016; Tsang et al., 2014; Mirdamadi and Block, 2021). Additionally, a sham cTBS protocol was implemented over S1, with the coil oriented sideways on the scalp.

### Transcutaneous electrical nerve stimulation (TENS)

TENS was applied to the median nerve at the right wrist using a bar electrode. A constant-current stimulator (DS7R, Digitimer Ltd.) delivered five square-wave pulses, each with a duration of 1 ms at a frequency of 10 Hz. This pulse train, consisting of five pulses over a span of 0.5 s, was followed by an equivalent period of 0.5 s without stimulation, maintaining a 50% duty cycle (Kaelin-Lang et al., 2002; Veldman et al., 2018). This sequence of stimulation was continuously applied for a total duration of 20 minutes. The intensity of the stimulus was calibrated to be below the motor threshold. This threshold was individually determined as the lowest intensity required to elicit observable muscle contractions. The low stimulation intensity and the 1 ms stimulus duration primarily activate large cutaneous and proprioceptive sensory fibers (Panizza et al., 1992). In the sham TENS protocol, the electrode configuration remained the same; however, the actual stimulation was applied for only 30 seconds.

### Study Design

This study comprised three distinct experiments. Specifically, Experiment 1 was designed to investigate the effects of cTBS applied to S1 and TENS of the median nerve on corticospinal excitability, as measured by MEP before and at 10, 30, and 50 minutes following cTBS and TENS. Experiment 2 aimed to examine the effects of cTBS applied to S1 and TENS of the median nerve on distinct intracortical mechanisms in M1, assessed by SICI and ICF. Experiment 3 focused on evaluating the effects of cTBS applied to S1 and TENS of the median nerve on S1-M1 interactions, as measured by SAI.

### Experiment 1

A total of fifty-two participants were enrolled in Experiment 1 and were randomly assigned to either the cTBS (n=26) or TENS (n=26) procedures. Within the cTBS procedure, participants were further divided into active cTBS (n=13) and sham cTBS (n=13) groups. Similarly, within the TENS procedure, participants were allocated to either the active TENS (n=13) or sham TENS (n=13) groups. For MEP measurements conducted before and after cTBS and TENS procedures, the stimulus intensity was set to evoke MEPs greater than 1 mV in at least 5 out of 10 consecutive trials. The specific stimulus intensity, predetermined for each participant, was consistently applied to induce MEPs at various time points, both pre- and post-procedures. Measurements were recorded at four intervals: 15 MEPs at baseline (5 minutes prior to the procedures), 15 MEPs at 10 minutes post-procedures, 15 MEPs at 30 minutes post-procedures, and 15 MEPs at 50 minutes post-procedures, totaling sixty MEPs. The inter-trial intervals for the MEP measurements had a median duration of 8 s, with a range from 7.2 to 8.8 s.

### Experiment 2

A total of fifty-two participants were recruited for Experiment 2. They were randomly allocated to either the cTBS procedure (n=26) or the TENS procedure (n=26). In both the cTBS and TENS procedures, participants received either active or sham stimulation. SICI and ICF were measured before and after the cTBS and TENS procedures. The measurements of SICI and ICF were randomized and separated by a 5-minute interval. For SICI measurements, the conditioning stimulus (CS) intensity was set at 70% of the RMT, while the test stimulus (TS) was administered at 120% of the RMT. A 2 ms inter-stimulus interval (ISI) was selected for SICI assessments. SICI was quantified by comparing the conditioned MEP size as a percentage of the unconditioned test MEP size. Fifteen conditioned and fifteen unconditioned MEPs were recorded both before and after cTBS and TENS procedures, resulting in a total of sixty MEPs per participant. For ICF measurements, the CS was delivered at 80% of the RMT, with the TS set at 120% of the RMT. The CS was applied 10 ms prior to the TS. To calculate ICF, the conditioned MEP size was measured as a percentage of the unconditioned test MEP size. Similarly, fifteen conditioned and fifteen unconditioned MEPs were recorded before and after cTBS and TENS procedures, totaling sixty MEPs per participant.

### Experiment 3

In Experiment 3, fifty-two participants were randomly assigned to either the cTBS procedure (n=26) or the TENS procedure (n=26). In each procedure, participants received either active or sham stimulation. For SAI measurements conducted before and after cTBS and TENS procedures, the CS was applied to the median nerve at an intensity equivalent to the peripheral motor threshold, with a pulse duration of 0.1 ms. This CS preceded the TS by an ISI of 20 ms. SAI was quantified by comparing the size of the MEP elicited by the conditioned stimulus to that of the unconditioned test MEP, expressed as a percentage. For each participant, we recorded fifteen conditioned and fifteen unconditioned MEPs both before and after cTBS and TENS procedures, resulting in a total of sixty MEPs per participant.

### Data Analysis

The Shapiro-Wilk test was utilized to assess the normality of distributions, while Levene’s test for equality of variances and Mauchly’s test of sphericity were employed to evaluate variance homogeneity. When assumptions of normal distribution were violated, data underwent logarithmic transformation. The Greenhouse-Geisser correction was applied if sphericity could not be assumed. In Experiment 1, mixed ANOVAs were used to examine the impact of various procedures (cTBS, sham cTBS, TENS, and sham TENS) and different time intervals (baseline, 10, 30, and 50 minutes) on MEP amplitude. In Experiment 2, mixed ANOVAs assessed the effects of the procedures (cTBS, sham cTBS, TENS, and sham TENS) and time (pre and post) on SICI and ICF. In Experiment 3, mixed ANOVAs analyzed the effects of the procedures (cTBS, sham cTBS, TENS, and sham TENS) and time (pre and post) on SAI. Statistical significance was set at p < 0.05. Group data are presented as means ± SD in the text.

## Results

### Cortical and peripheral somatosensory stimulation enhance corticospinal excitability

Figure 1A illustrates the changes in MEP amplitude for a representative participant at baseline, and at 10, 30, and 50 minutes post-stimulation during S1 stimulation via cTBS. The MEP amplitude increased at all time intervals following cTBS compared to baseline measurements. Figure 2A depicts the alterations in MEP amplitude for a representative participant at baseline, and at 10, 30, and 50 minutes post-stimulation during peripheral stimulation via TENS. An increase in MEP amplitude was observed at 10 minutes post-TENS, although no changes were detected at 30 and 50 minutes when compared to baseline measurements. A mixed ANOVA analysis revealed significant interaction effects between different procedures and time intervals on MEP amplitude (F(9, 144)=2.317, p=0.018). Post hoc comparisons revealed that after S1 stimulation via cTBS, MEP amplitude demonstrated a significant increase at 10 minutes (1.47±0.96 mV; p=0.008), 30 minutes (1.61±1.10 mV; p=0.011), and 50 minutes (1.63±1.07 mV; p=0.010) compared to baseline (1.04±0.56 mV; Figure 1B). Conversely, following sham cTBS, the MEP amplitude remained relatively unchanged at 10 minutes (0.78±0.33 mV; p=0.1), 30 minutes (0.88±0.50 mV; p=0.4), and 50 minutes (0.90±0.54 mV; p=0.5) compared to baseline (0.90±0.42 mV; Figure 1C). Following peripheral stimulation via TENS, post hoc comparisons indicated a significant increase in MEP amplitude at 10 minutes (1.54±0.52 mV; p=0.006), but no significant changes were observed at 30 minutes (1.33±0.42 mV; p=0.2) or 50 minutes (1.35±0.61 mV; p=0.2), compared to baseline (1.24±0.38 mV; Figure 2B). After sham TENS, MEP amplitude remained relatively stable at 10 minutes (1.42±1.13 mV; p=0.3), 30 minutes (1.40±1.20 mV; p=0.2; Figure 1C), and 50 minutes (1.30±0.96 mV; p=0.1) compared to baseline (1.54±1.02 mV; Figure 2C). Figures 1D and 2D display group data comparing changes in MEP amplitude, expressed as MEP amplitudes normalized to baseline, at various intervals between active stimulation and sham stimulation during the cTBS and TENS procedures. A mixed ANOVA analysis revealed significant interaction effects between different procedures and time intervals on normalized MEP amplitude (F(9, 144)=2.684, p=0.007). Post hoc comparisons indicated that normalized MEP amplitude significantly increased in cTBS compared to sham cTBS at 10 minutes (p=0.003), 30 minutes (p=0.01), and 50 minutes (p=0.004). Normalized MEP amplitude also significantly increased in TENS compared to sham TENS at 10 minutes (p=0.01), but not at 30 minutes (p=0.1) or 50 minutes (p=0.1). Collectively, these results demonstrate that both S1 and peripheral stimulations via cTBS and TENS influence corticospinal excitability; however, the effect of S1 stimulation is more enduring, lasting up to 50 minutes, compared to the brief, 10- minute impact of peripheral stimulation.

**Figure 1.**
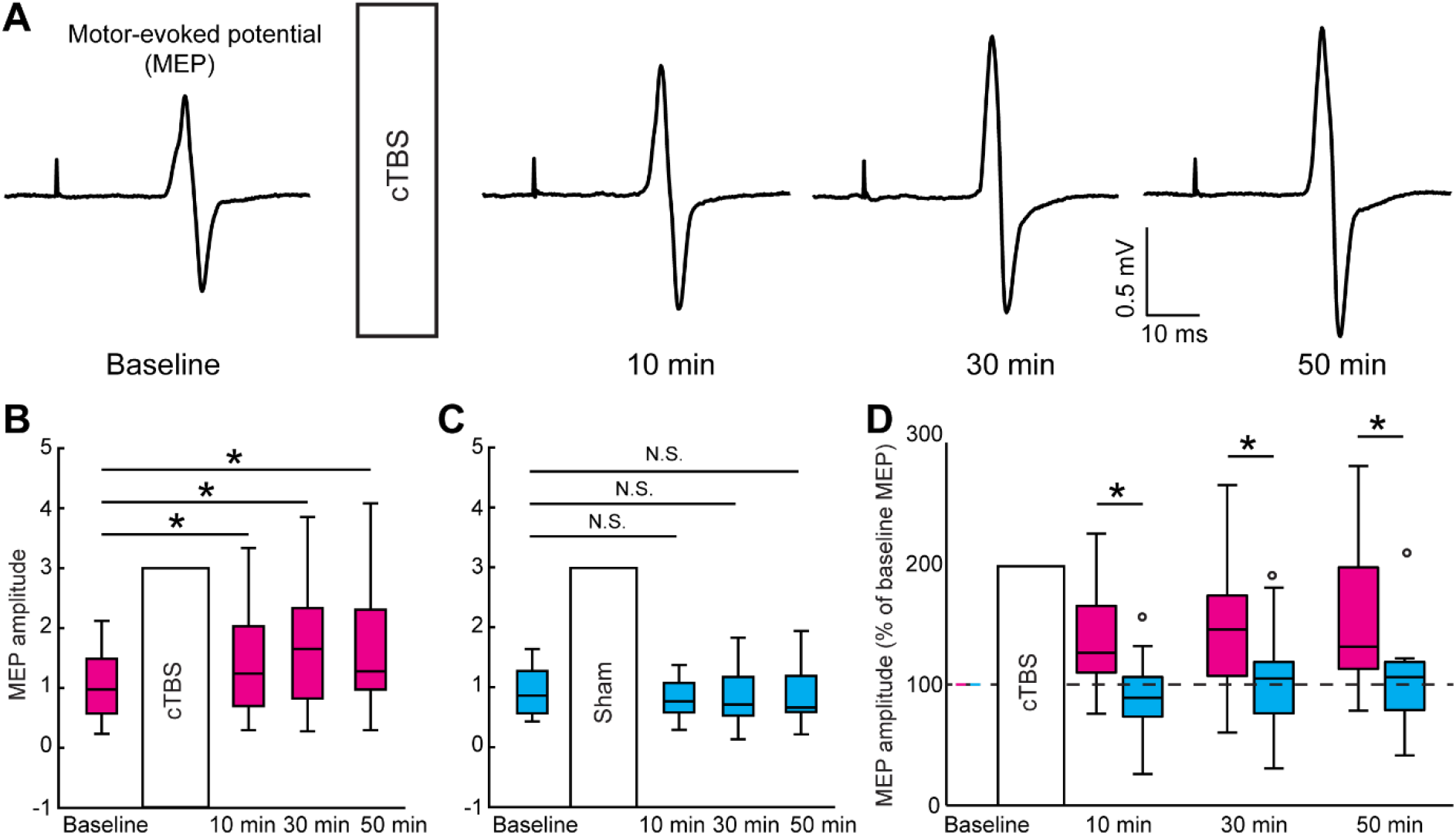
Corticospinal Excitability Modulated by cTBS. (A) MEP waveforms from a representative subject recorded at baseline and 10, 30, and 50 minutes after the cTBS intervention. (B) Boxplot showing the MEP amplitudes across time points (baseline and 10, 30, and 50 minutes after the cTBS intervention). (C) Boxplot illustrating the MEP amplitudes throughout the sham cTBS intervention procedure. (D) Comparison of normalized MEP amplitudes between the cTBS and sham cTBS interventions. * p < 0.05; N.S. p >=0.05; ° outlier.

**Figure 2.**
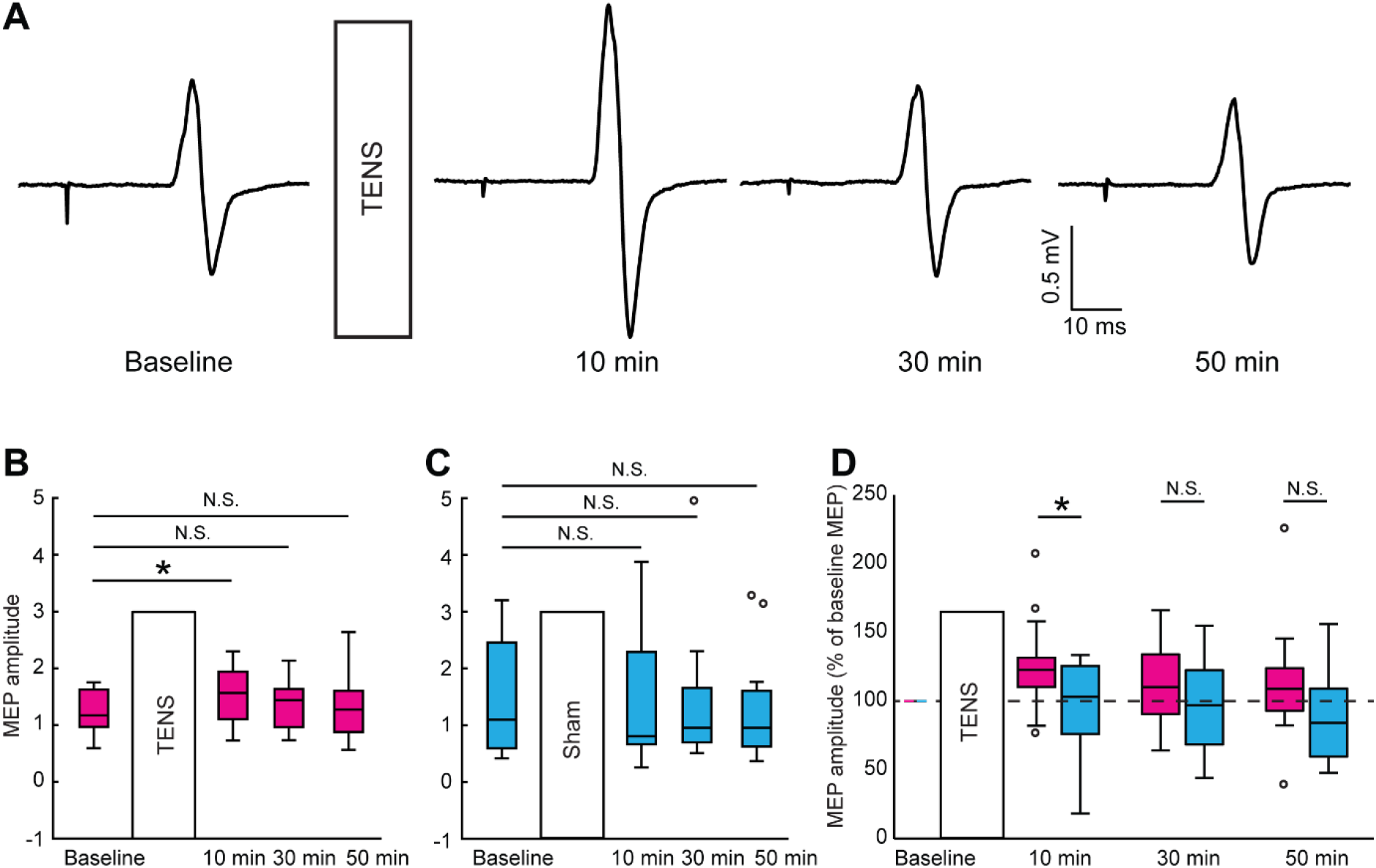
Corticospinal Excitability Modulated by TENS. (A) MEP waveforms from a representative subject recorded at baseline and 10, 30, and 50 minutes after the TENS intervention. (B) Boxplot showing the MEP amplitudes across time points (baseline and 10, 30, and 50 minutes after the TENS intervention). (C) Boxplot illustrating the MEP amplitudes throughout the sham TENS intervention procedure. (D) Comparison of normalized MEP amplitudes between the TENS and sham TENS interventions. * p < 0.05; N.S. p >=0.05; ° outlier.

### Effects of cortical and peripheral somatosensory stimulation on SICI, ICF, and SAI

Figures 3A and 3C illustrate the SICI assessment conducted before and after the administration of cTBS and TENS, respectively, with the test MEP shown in black and the conditioned MEP shown in red. When a subthreshold conditioned TMS was applied to M1 2 ms prior to the test TMS over M1, the conditioned MEP demonstrated significant inhibition compared to the test MEP. To confirm the presence of SICI prior to the procedures (cTBS, sham cTBS, TENS, and sham TENS), paired t-tests were conducted to compare conditioned MEPs with test MEPs before each procedure. The analysis indicated a significant decrease in the amplitudes of conditioned MEPs relative to the amplitudes of test MEPs across all procedures, including cTBS (p<0.001), sham cTBS (p<0.001), TENS (p<0.001), and sham TENS (p<0.001). These results confirmed SICI’s occurrence before each of the four interventions. Figures 3B and 3D indicate that the degree of SICI observed prior to the administration of cTBS and TENS was comparable to the extent of SICI recorded following these procedures. A mixed ANOVA analysis revealed no significant effects of the procedures (F(3, 48)=0.299, p=0.826), timing (F(1, 48)=1.157, p=0.287), or their interaction (F(3, 48)=0.103, p=0.958) on SICI. The SICI values were similar before and after cTBS (pre-cTBS=45±18%; post-cTBS=42±19%), sham cTBS (pre-sham cTBS=47±24%; post-sham cTBS=47±26%), TENS (pre-TENS=43±19%; post-TENS =41±14%), and sham TENS (pre-sham TENS=42±16%; pre-sham TENS=40±14%). These results indicate that cortical and peripheral somatosensory stimulation do not affect SICI.

**Figure 3.**
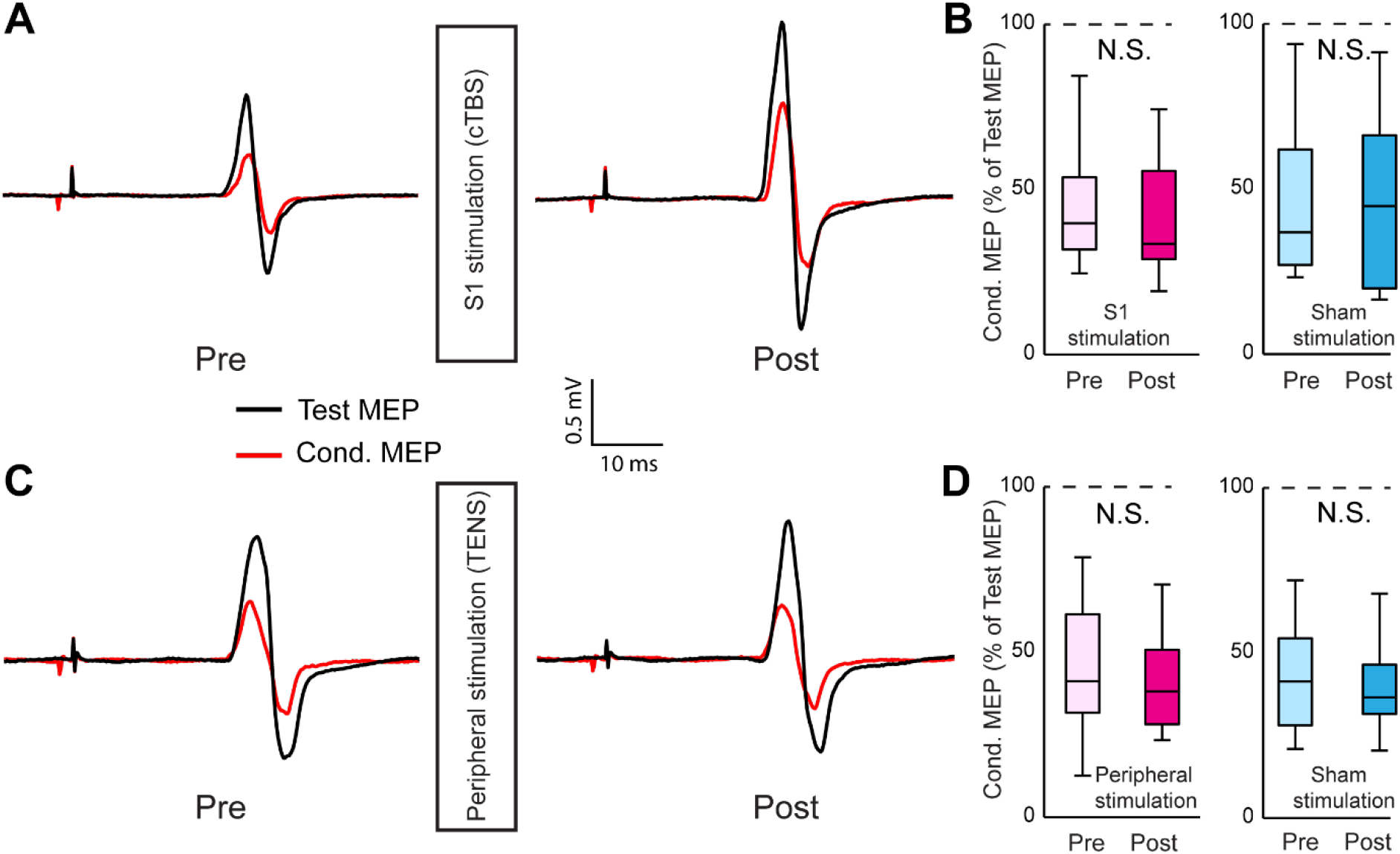
(A) Representative examples of MEPs induced by single- and paired-pulse (ISI = 2 ms) TMS, demonstrating short-latency intracortical inhibition (SICI) before and after the cTBS intervention. (B) Boxplots showing the effects of cTBS and sham cTBS on the magnitudes of conditioned MEPs, expressed as % of test MEP. (C) Representative examples of MEPs induced by single- and paired-pulse TMS, demonstrating SICI across the TENS and sham TENS procedures. (D) Boxplots showing SICI before and after the TENS (or sham TENS) intervention. N.S. p >=0.05.

Figures 4A and 4C depict the ICF assessment conducted before and after the administration of cTBS and TENS, respectively. The test MEP is shown in black, while the conditioned MEP is presented in red. It is noteworthy that the conditioned MEP demonstrated facilitation compared to the test MEP when a subthreshold conditioned TMS was applied to M1 10 ms prior to the test TMS over M1. To confirm the presence of ICF prior to the procedures (cTBS, sham cTBS, TENS, and sham TENS), paired t-tests were performed. The results indicated an increase in conditioned MEP amplitudes relative to test MEPs for all procedures, with significant p-values (p<0.016). This established the occurrence of ICF before each procedure. Figures 4B and 4D demonstrate that the degree of ICF observed prior to the administration of cTBS and TENS was comparable to the ICF recorded following these procedures. The mixed ANOVA for the ICF measurement revealed no significant intervention effects (F(3, 48)=0.132, p=0.941), timing effects (F(1, 48)=1.382, p=0.246), or interaction effects (F(3, 48)=0.167, p=0.918). The ICF values remained consistent before and after cTBS (pre-cTBS=139±50%; post-cTBS=133±46%), sham cTBS (pre-sham cTBS=134±33%; post-sham cTBS=126±30%), TENS (pre-TENS=134±43%; post-TENS =133±50%), and sham TENS (pre-sham TENS=132±35%; post-sham TENS=123±25%). These results indicate that neither cortical nor peripheral somatosensory stimulation affects ICF.

**Figure 4.**
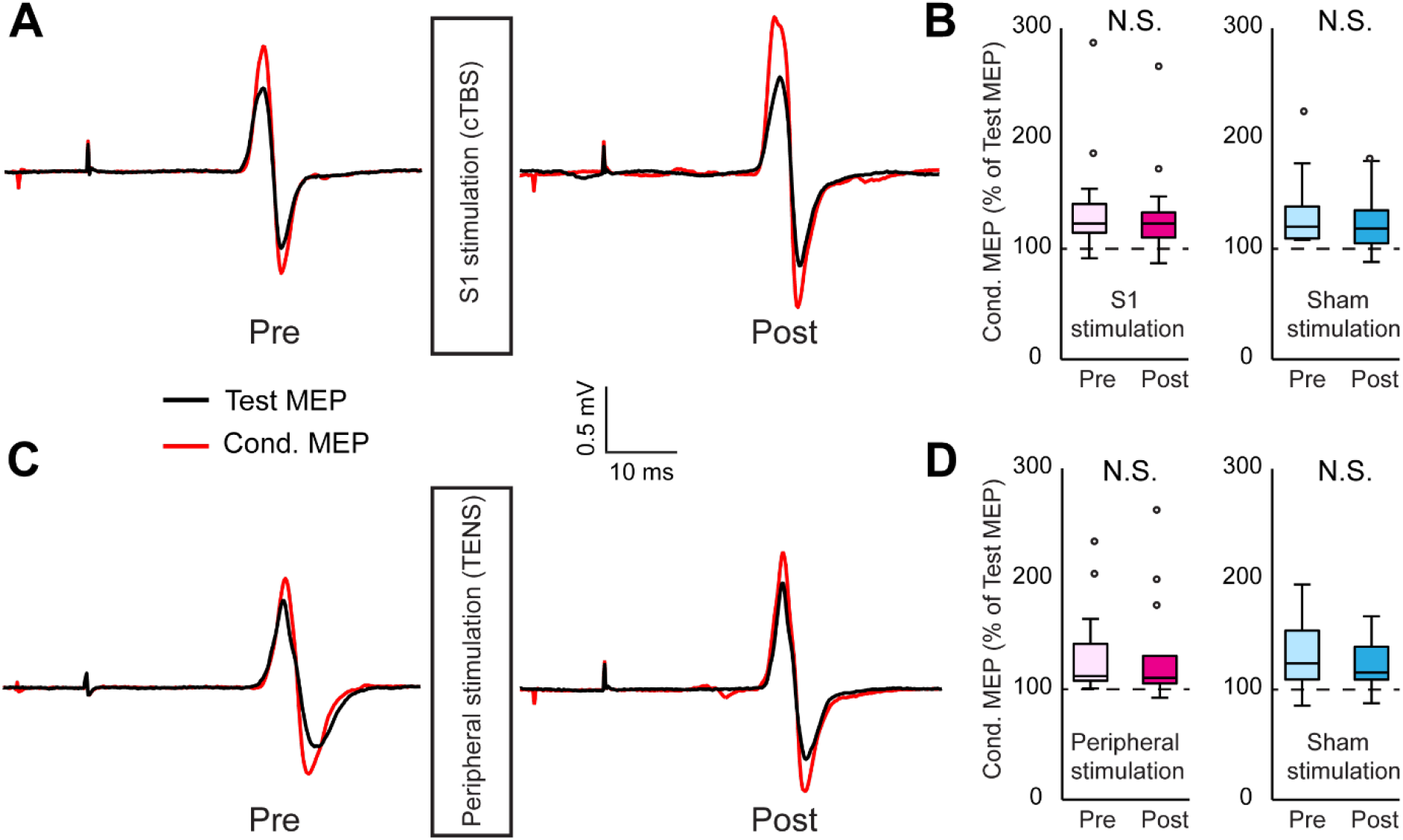
(A) Representative examples of MEPs induced by single- and paired-pulse (ISI = 10 ms) TMS, demonstrating intracortical facilitation (ICF) before and after the cTBS intervention. (B) Boxplots showing the effects of cTBS and sham cTBS on the magnitudes of conditioned MEPs, expressed as % of test MEP. (C) Representative examples MEPs induced by single- and paired-pulse TMS, demonstrating ICF across the TENS and sham TENS procedures. (D) Boxplots showing ICF before and after the TENS (or sham TENS) intervention. N.S. p >=0.05; ° outlier.

Figures 5A and 5C illustrate the SAI assessment conducted before and after the administration of cTBS and TENS, respectively. The test MEP is displayed in black, while the conditioned MEP is shown in red. Notably, the conditioned MEP exhibited inhibition compared to the test MEP when electrical stimulation was applied to the median nerve at an intensity equivalent to the peripheral motor threshold 20 ms prior to the test TMS over M1. To confirm the presence of SAI prior to the procedures (cTBS, sham cTBS, TENS, and sham TENS), paired t-tests were utilized. Results indicated a significant decrease in conditioned MEP amplitudes, confirming SAI occurrence before each intervention (p<0.001). The results for the SAI measurement demonstrated significant effects from timing (F(1, 48)=6.202, p=0.016) and marginally significant effects from the interaction between procedures and timing (F(3, 48)=2.564, p=0.066), but not from the procedures alone (F(1, 48)=1.372, p=0.262). Post-hoc analysis revealed that SAI changed from 63±24% pre-stimulation to 86±36% post-stimulation following cTBS (p=0.003; Figure 5B). Conversely, no significant change in SAI was observed during TENS before and after the stimulation (pre-TENS=55±23%; post- TENS =59±27%; p=0.2). Additionally, no significant differences in SAI were found before and after either sham cTBS (pre-sham cTBS=59±28%; post-sham cTBS=65±23%; p=0.2) or sham TENS (pre-sham TENS=60±22%; post-sham TENS=59±28%; p=0.5). These findings suggest that cortical somatosensory stimulation modulates SAI, whereas peripheral stimulation does not affect SAI.

**Figure 5.**
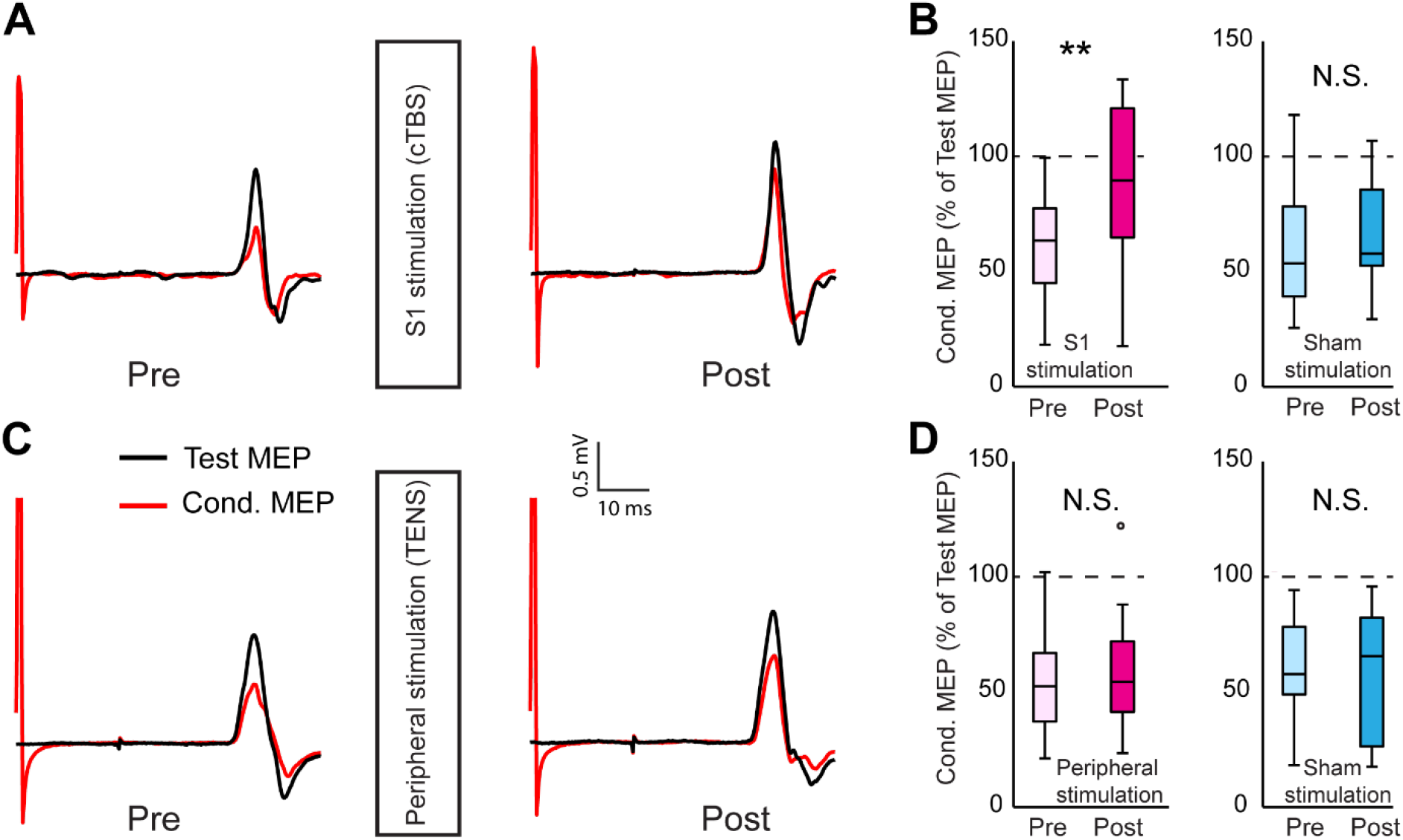
(A) Representative examples of MEPs induced by single- and paired-pulsed (ISI = 20 ms) stimulation, demonstrating short-latency afferent inhibition (SAI) before and after the cTBS intervention. (B) Boxplots showing the effects of cTBS and sham cTBS on the magnitudes of conditioned MEPs, expressed as % of test MEP. (C) Representative examples MEPs induced by single- and paired-pulse stimulation, demonstrating SCI across the TENS and sham TENS procedures. (D) Boxplots showing SAI before and after the TENS (or sham TENS) intervention. * p < 0.05; N.S. p >=0.05; o outlier.

## Discussion

This study investigated the distinct effects of cortical and peripheral somatosensory stimulation on M1 plasticity. cTBS was applied to S1, while TENS was administered to the median nerve. The subsequent impacts on corticospinal excitability, intracortical circuits within M1, and interactions between S1 and M1 were measured. The results showed that both cTBS and TENS increased MEP amplitude, indicating enhanced corticospinal excitability. However, the effect induced by cTBS was significantly more prolonged, lasting 50 minutes compared to 10 minutes for TENS. Neither cTBS nor TENS affected short-interval intracortical inhibition (SICI) or intracortical facilitation (ICF), suggesting no impact on GABAergic or glutamatergic intracortical circuits within M1. Notably, cTBS, but not TENS, reduced short-afferent inhibition (SAI), indicating specific modulation of S1-M1 interactions. Sham stimulation did not produce significant changes in any measured parameter. These results elucidate distinct mechanisms through which cortical and peripheral somatosensory stimulation influence M1 plasticity, highlighting the potential for targeted interventions to modulate motor function and inform the development of therapeutic strategies.

Our first experiment demonstrated that both cTBS applied to S1 and TENS of the median nerve resulted in an increased MEP amplitude. This finding supports the hypothesis that cortical and peripheral somatosensory stimulation can modulate corticospinal excitability, corroborating previous studies which have indicated that such stimulations lead to heightened corticospinal excitability (Ridding et al. 2000; Kaelin-Lang et al., 2002; Veldman et al., 2018; Jacobs et al., 2012, 2014). The mechanism for corticospinal facilitation following cortical somatosensory stimulation may involve a direct cortico-cortical pathway from S1 to M1. Dense anatomical connections exist between S1 and M1 (Donoghue and Parham, 1983; Mao et al., 2011; Veinante and Deschênes, 2003), enabling information to be conveyed through cortico-cortical pathways between these regions (Guo et al., 2018; Suresh et al., 2020; Umeda et al., 2019; Brown et al., 2019; Davis et al., 2022). Electrophysiological studies have documented that activity in M1 reflects activity in S1 during movements (Umeda et al., 2019), and intra-cortical stimulation of S1 generates evoked responses in M1 (Osborn et al., 2021). Further studies have shown that MEP size was reduced when the ipsilateral S1 was stimulated 1ms prior to M1 at a supra-threshold stimulation intensity (Brown et al., 2019; Davis et al., 2022), suggesting an inhibitory pathway from S1 to M1 (S1-M1 inhibition). Evidence has shown that theta-burst TMS can produce long-lasting alterations in neural excitability thought to involve long-term depression (LTD)- or long-term potentiation (LTP)-like plasticity (Huang et al., 2005; Suppa et al., 2016; Rounis and Huang, 2020). Specifically, cTBS decreases S1 excitability (LTD-like plasticity; Jacobs et al., 2012, 2014; Kumar et al., 2019; Wang et al., 2024). Therefore, cTBS applied to S1 may act through a cortico-cortical route to potentiate M1 excitability via an inhibitory pathway from S1 to M1, leading to an increase in corticospinal output neurons in M1. The mechanisms underlying corticospinal facilitation following peripheral somatosensory stimulation remain largely unexplored, but they likely involve the interplay between S1 and M1. TENS applied to the median nerve activates both muscle and cutaneous afferents, which project to S1 and subsequently connect to M1 (Friedman and Jones, 1981; Jones, 2000). The most probable pathway mediating changes in corticospinal excitability involves sensory information relayed from peripheral receptors to M1 via S1, specifically through the thalamo-cortical pathway. This is supported by anatomical studies demonstrating dense fiber tracts connecting the thalamus to both S1 and M1 (Cerkevich et al., 2014; Cheema et al., 1985; Hatanaka et al., 2005). S1, receiving somatosensory input from the thalamus, possesses extensive reciprocal connections with M1 (Ghosh et al., 1987; Kaas et al., 1983; Keysers et al., 2010), further supporting its role as a relay station. Notably, previous research has shown that TENS does not affect peripheral nerve conduction, as evidenced by the lack of changes in maximal compound action potential, F-wave, or responses to brainstem stimulation (Hamdy et al., 1998; Kaelin-Lang et al., 2002; Ridding et al., 2000; Stefan et al., 2000). This suggests that the observed increases in corticospinal excitability after TENS are not due to changes at the neuromuscular junction or peripheral nerve level but rather occur at a supraspinal level, most likely involving S1-M1 interactions via the thalamo-cortical pathway.

The significant difference in effect duration—50 minutes for cTBS compared to 10 minutes for TENS—suggests distinct underlying mechanisms of action. The prolonged increase in MEP amplitude following cTBS is likely attributable to its direct modulation of S1 excitability. cTBS is thought to induce LTD-like plasticity, resulting in sustained changes in the excitability of cortico-cortical projections from S1 to M1. Conversely, the transient increase in MEP amplitude observed after TENS likely reflects a different mechanism. Peripherally applied TENS influences afferent input to the cortex, potentially leading to a temporary increase in the synaptic efficacy of sensory afferents projecting to S1, which subsequently modulates M1 activity. However, this effect appears to be short-lived, indicating that the underlying changes in synaptic strength are not as enduring as those induced by direct cortical stimulation with cTBS. This difference in the duration of effects underscores the distinct ways in which cortical and peripheral somatosensory stimulation interact with and modulate M1 circuitry.

Our study revealed that neither cTBS applied to S1 nor TENS of the median nerve altered GABA-mediated short-interval intracortical inhibition (SICI) or glutamate-mediated intracortical facilitation (ICF). This finding aligns with previous research indicating that somatosensory stimulation does not modulate these intracortical circuits (Celnik et al., 2007; Conforto et al., 2010; Kaelin-Lang et al., 2002; Sawaki et al., 2006; Jacobs et al., 2014). Therefore, it appears that cortical and peripheral somatosensory stimulation do not affect the interneurons associated with SICI and ICF. Instead, the observed changes in corticospinal excitability likely result from other mechanisms, such as direct S1-M1 projections or modulation of sensory afferent input along ascending pathways. Furthermore, we observed a differential effect of cTBS and TENS on SAI. Specifically, cTBS applied to S1 significantly reduced SAI compared to baseline. We propose that cTBS over S1 diminishes S1 excitability, consequently reducing its modulatory influence on M1 and leading to the observed decrease in SAI. This interpretation is consistent with previous studies demonstrating that cTBS over S1 decreases high-frequency oscillations within S1 (Ishikawa et al., 2007), impairs tactile acuity (Rai et al., 2012), and diminishes SAI (Tsang et al., 2014; Bao et al., 2024). Conversely, TENS did not influence SAI, indicating that TENS may not alter S1 excitability and therefore does not affect S1’s modulatory effect on M1, as indicated by SAI. This is supported by studies showing unchanged somatosensory evoked potentials after TENS administration (Kaelin-Lang et al., 2002).

This study acknowledges several limitations. First, the impact of varying cTBS and TENS parameters (such as duration, intensity, and frequency) on M1 plasticity was not investigated. These parameters can influence the effects of somatosensory stimulation. For instance, Goldsworthy et al. (2012) found that 50 Hz cTBS initially suppressed MEPs, whereas 30 Hz cTBS led to sustained suppression. Similarly, the effect of TENS on MEP amplitude fluctuates based on frequency and duration (Chipchase et al., 2011; Veldman et al., 2015; Sato et al., 2022). Future research should systematically investigate these parameters to optimize cTBS and TENS for modulating M1 plasticity. Second, only the short-term effects of cTBS and TENS were assessed in this study. Longitudinal studies are required to ascertain the long-term impact of these interventions on M1 plasticity and to evaluate the potential cumulative benefits of repeated cortical and peripheral somatosensory stimulation. Third, although sham stimulation was included, it may not perfectly replicate real cTBS or TENS, which could affect participant blinding and introduce a placebo effect. Future studies should explore alternative sham procedures to address this limitation.

In conclusion, this study demonstrates that both cortical (cTBS) and peripheral (TENS) somatosensory stimulation can enhance corticospinal excitability, as evidenced by increased MEP amplitudes. However, these two modalities exert their effects through distinct mechanisms. cTBS induces a prolonged increase in M1 excitability, likely through long-lasting modulation of cortico-cortical pathways between S1 and M1, including a reduction in S1-M1 inhibitory interactions (as evidenced by decreased SAI). In contrast, TENS produces a more transient increase in M1 excitability, potentially by temporarily enhancing the synaptic efficacy of sensory afferents projecting to S1. Neither intervention affected intracortical circuits within M1, as reflected by unchanged SICI and ICF measures. These findings highlight the complexity of the interactions between the somatosensory and motor systems and suggest that targeted somatosensory stimulation holds promise for modulating motor function and developing novel therapeutic strategies. While our findings contribute significantly to understanding the distinct effects of cTBS and TENS, further research is needed to address the limitations of this study, including optimizing stimulation parameters, investigating long-term effects, and exploring the clinical applicability of these techniques in populations with neurological disorders.

## Data availability statement

The data that support the findings of this study are available from the corresponding author upon reasonable request.

